# Elucidation of novel miRNA candidates and their role in unraveling the pathology of Non-Alcoholic Fatty Liver Disease

**DOI:** 10.1101/2020.01.31.917831

**Authors:** Bincy Mary Biju, Saheb Singh Chhabra, Chiradeep Sarkar

## Abstract

Non-Alcoholic Fatty Liver Disease (NAFLD) is a chronic liver disease which is observed in people who do not abuse alcohol. Main cause of NAFLD is Non-Alcoholic Steatohepatitis (NASH) where there is accumulation of fats such as triglyceride in liver and the disease progression ranges from simple steatosis to fibrosis and cirrhosis. MicroRNAs are critical players in post-transcriptional gene regulation of diseases with complex etiology. In this study, we have elucidated the role of microRNAs (miRNAs) in the pathophysiology of NAFLD/NASH and unravelled molecular markers for diagnosis of NAFLD. A subset of genes (n=10) responsible for NAFLD/NASH were selected and detailed *in silico* analysis carried out using multiple tools. miRDB and DIANA-microT were used to find putative miRNA binding sites followed by analysis using miRTarBase which is an experimentally validated database of miRNA-target interactions. The study elucidated a number of statistically significant predictions for both miRDB (scores >80) and DIANA-microT (values >0.90) and also strong experimental validation in miRTarBase. The analysis revealed that certain miRNAs like miR-7 & miR-548 family members are found in both the programmes, miRDB and DIANA-microT, targeting genes involved in liver function. They were also identified in the experimental validation database miRTarBase. These miRNAs probably play an important role in the pathophysiology of this disease. They can also be used as prognostic/diagnostic markers for assessment of NAFLD.

## Introduction

Non-alcoholic fatty liver disease or NAFLD is a chronic liver disease which leads to liver damage. This disease is seen in individuals who do not abuse alcoholic substances but still have the similar damage as an alcohol-induced damaged liver. Accumulation of different fats in liver cells is the reasons for NAFLD. This accumulation is caused due to metabolic syndrome and obesity.^1^ As per population-based screening, at least 25% of US population suffers from NAFLD ^2^ and study conducted by Ankita *et.al,* in 2015, the prevalence of NAFLD in India is 9% in rural population and 32% in urban population.^3^

The individual suffering from NAFLD tends to develop NASH (a subset disease to NAFLD). There are no non-invasive tests to determine and differentiate between NAFLD and NASH which makes it more difficult to understand the exact incidence and prevalence of these diseases. However, use of serum aminotransferases as a surrogate marker for NASH, by a population-based study suggests that, about 6-8% of the adults suffering from NAFLD has NASH (that is; 25% of the population suffering from NAFLD).^4^ Small liver biopsy study shows that 25% of patients with NASH develop cirrhosis as their condition progresses.^5^ Considering these studies as representative to larger population, about 1-2.5% of the US population suffers from cirrhosis due to NAFLD/NASH. Cirrhosis is a leading factor for liver cancer in humans. However, growing number of liver cancer patients shows that liver cancer has been developing in individuals with NAFLD even though they don’t suffer from cirrhosis. This shows that the risk factors for NAFLD like obesity and metabolic syndrome are also responsible for development of cancer in extra hepatic tissue.^6^

Patients with NAFLD are a higher risk of developing hepatocellular carcinoma (HCC). Out of all the HCC cases in Western countries, 4-22% of the cases are caused due to NAFLD. ^7^ NAFLD affecting 30% of the general population in North America and is closely associated with the growing obesity problem that people suffer from. Among this population, it is seen that 30-50% of the people suffer from NASH (Non-Alcoholic Steatohepatitis). Studies show that about 70% of the patients with NAFLD have normal liver enzymes.^8^ However, alanine aminotransferases (ALT) appears to be seven-fold higher in obese people.^9^ Some people with normal aminotransferase have been seen with NASH and indicative fibrosis. Thus, these surveys confirm, that the entire histopathological band of NAFLD could be seen in people with normal liver enzymes.^10^ In Asia, where viral hepatitis is still an epidemic, 1-2% of the HCC cases are because of NAFLD.^11^ Globally, cryptogenic cirrhosis is responsible for 15-30% of HCC cases and about half of these are believed to have risen due to NAFLD.^12^

miRNAs or microRNAs, as the name suggests are small, non-coding RNA molecules which are involved in regulating gene expression at post-transcriptional level. The miRNAs are around 19-23 nucleotides long. MicroRNA was discovered by Ambrose and colleagues in 1993.^13^ They regulate the gene expression by interfering in translation process.^14,15^ From multiple studies, it is evident now that microRNA plays an important role in pathophysiology of complex multi-factorial disorders including NAFLD/NASH.^14,16,17^ Studies have shown the role of miR-122 in progression of severe liver injury such as fibrosis.^18^ Liver-specific knockout animal models have shown development and fibrosis, and Steatohepatitis.^18^ miR-34a is involved in lipid metabolism, apoptosis, cell cycle control and studies in mice have shown that overexpression of miR-34a increases hepatic Triglyceride (TG) level. ^19^ Studies have shown that elevated miR-103 cause’s stimulation of ectopic lipid accumulation in liver causing NAFLD. ^20,21,22^ Studies on NAFLD patients have also that down regulation of miR-155 enhances LXRα/SREBP-1c signal which lead to accumulation of lipids in liver.^23^ Animal studies have shown that inhibition miR-29a causes enhancement of lipoprotein lipase (LPL) expression thereby causing lipid accumulation in liver. ^24^ Studies on humans showed that, up regulation of miR-192 have co-relation with liver disease.^14^ However, we still don’t know the involvement of various miRNAs and their role in the pathophysiology of NAFLD/NASH. Hence, a subset of genes responsible for NAFLD/NASH was selected from the study of Alessandra *et. al.*, and a detailed *in silico* work was carried out in this present study.^16^ It is hoped that this information would serve as the basis for further studies in elucidating the role of microRNAs in the pathogenesis of NAFLD/NASH as well as provide diagnosis / prognostic markers for NAFLD/NASH.

## Methodology

The input gene dataset for this study included candidate genes from a spectrum of genetic factors implicated in the complex etiology of NAFLD. This present list of genes and their role and importance in NAFLD has been discussed in detail in the paper of Alessandra *et. al.* ^16^. In all, 10 genes different physiological categories were selected for identifying the various miRNAs targeting these genes. The list of these 10 candidate genes and their gene description are presented in Table 1 of this study.

**Table 1:** The list of 10 candidate genes implicated in the complex etiology of NAFLD and used as input for miRNA elucidation pipeline.

| Gene name | Gene Description |
| --- | --- |
| APOB | apolipoprotein B |
| ENPP1 | ectonucleotide pyrophosphatase/phosphodiesterase 1 |
| GCKR | glucokinase (hexokinase 4) regulator |
| HFE | hemochromatosis |
| PNPLA3 | phospholipase domain containing 3 |
| MBOAT7 | membrane bound O-acyltransferase domain containing 7 |
| IRS1 | insulin receptor substrate 1 |
| SLC2A1 | solute carrier family 2 (facilitated glucose transporter), member 1 |
| TM6SF2 | transmembrane 6 superfamily member 2 |
| TMC4 | transmembrane channel-like 4 |

### miRDB analysis

For the task of identifying the various miRNAs targeting these different genes involved in NAFLD pathophysiology, the miRDB database was used.^25^ miRDB is a database for miRNA target prediction and annotation. The various genes selected in this study were analysed by a tool called Mir Target v4 integrated in miRDB and which functions by analysing thousands of genes regulated by miRNAs with a Support Vector Machines learning framework. Common features involved in miRNA target binding have been identified and incorporated in this tool to predict miRNA targets in species like human, rat, mouse, dog and chicken. The Mir Target v4 prediction algorithm has been validated by independent experimental data for predicting a large number of miRNA down-regulated gene targets. In this study, the selected genes were searched individually in miRDB by entering the appropriate ‘Gene Symbol’ in the search box and using ‘Human’ as the option for organism. Now for each gene of interest and if predicted to be targeted by mature miRNAs, this results in a table where the multiple mature miRNAs targeting that gene are ranked according to a score assigned by the target prediction algorithm. Other details like miRNA sequence, 3’UTR sequence and gene description are also given. According to the scoring scheme in miRDB, the prediction scores for predicted targets range between 50-100. However, higher the score, the more is the confidence in the prediction and scores >80 are considered to be most likely to be real. Based on this, Table 2, was prepared having only those hits which have scores above or equal to 80 for the various genes predicted to be targeted by miRNAs.

**Table 2:** List of candidate genes with various miRNAs targeting them and with cut-off score ≥ 80 according to miRDB analysis (Mir Target v4).

| Gene symbol | Target score | miRNA name |
| --- | --- | --- |
| APOB | 94 | <u>hsa-miR-5008-3p</u> |
| APOB | 94 | <u>hsa-miR-7157-3p</u> |
| APOB | 94 | <u>hsa-miR-6737-3p</u> |
| APOB | 92 | <u>hsa-miR-6124</u> |
| APOB | 91 | <u>hsa-miR-449c-5p</u> |
| APOB | 91 | <u>hsa-miR-34b-5p</u> |
| APOB | 88 | <u>hsa-miR-548p</u> |
| APOB | 87 | <u>hsa-miR-2682-5p</u> |
| APOB | 87 | <u>hsa-miR-10522-5p</u> |
| APOB | 86 | <u>hsa-miR-1237-3p</u> |
| APOB | 84 | <u>hsa-miR-1468-3p</u> |
| APOB | 80 | <u>hsa-miR-7161-5p</u> |
| ENPP1 | 100 | <u>hsa-miR-3613-3p</u> |
| ENPP1 | 100 | <u>hsa-miR-3163</u> |
| ENPP1 | 98 | <u>hsa-miR-4424</u> |
| ENPP1 | 98 | <u>hsa-miR-371b-5p</u> |
| ENPP1 | 98 | <u>hsa-miR-616-5p</u> |
| ENPP1 | 98 | <u>hsa-miR-373-5p</u> |
| ENPP1 | 95 | <u>hsa-miR-548c-3p</u> |
| ENPP1 | 93 | <u>hsa-miR-607</u> |
| ENPP1 | 93 | <u>hsa-miR-4319</u> |
| ENPP1 | 92 | <u>hsa-miR-7110-3p</u> |
| ENPP1 | 92 | <u>hsa-miR-7854-3p</u> |
| ENPP1 | 91 | <u>hsa-miR-4307</u> |
| ENPP1 | 90 | <u>hsa-miR-3120-3p</u> |
| ENPP1 | 90 | <u>hsa-miR-6873-3p</u> |
| ENPP1 | 90 | <u>hsa-miR-125a-5p</u> |
| ENPP1 | 90 | <u>hsa-miR-125b-5p</u> |
| ENPP1 | 89 | <u>hsa-miR-1294</u> |
| ENPP1 | 89 | <u>hsa-miR-9986</u> |
| ENPP1 | 88 | <u>hsa-miR-7843-3p</u> |
| ENPP1 | 87 | <u>hsa-miR-202-5p</u> |
| ENPP1 | 86 | <u>hsa-miR-5586-5p</u> |
| ENPP1 | 85 | <u>hsa-miR-3169</u> |
| ENPP1 | 83 | <u>hsa-miR-556-3p</u> |
| ENPP1 | 83 | <u>hsa-miR-10523-5p</u> |
| ENPP1 | 83 | <u>hsa-miR-4316</u> |
| ENPP1 | 82 | <u>hsa-miR-6867-3p</u> |
| ENPP1 | 82 | <u>hsa-miR-6734-5p</u> |
| ENPP1 | 82 | <u>hsa-miR-4740-5p</u> |
| ENPP1 | 82 | <u>hsa-miR-4768-5p</u> |
| ENPP1 | 82 | <u>hsa-miR-338-5p</u> |
| ENPP1 | 82 | <u>hsa-miR-6857-5p</u> |
| ENPP1 | 82 | <u>hsa-miR-6833-3p</u> |
| ENPP1 | 80 | <u>hsa-miR-2117</u> |
| ENPP1 | 80 | <u>hsa-miR-12125</u> |
| ENPP1 | 80 | <u>hsa-miR-3200-5p</u> |
| GCKR | 83 | <u>hsa-miR-4306</u> |
| GCKR | 80 | <u>hsa-miR-4656</u> |
| HFE | 99 | <u>hsa-miR-5093</u> |
| HFE | 91 | <u>hsa-miR-5696</u> |
| HFE | 88 | <u>hsa-miR-361-5p</u> |
| HFE | 88 | <u>hsa-miR-1245b-3p</u> |
| HFE | 87 | <u>hsa-miR-7159-3p</u> |
| HFE | 86 | <u>hsa-miR-5700</u> |
| HFE | 85 | <u>hsa-miR-3915</u> |
| HFE | 83 | <u>hsa-miR-12124</u> |
| HFE | 82 | <u>hsa-miR-324-5p</u> |
| HFE | 82 | <u>hsa-miR-3662</u> |
| PNPLA3 | 90 | <u>hsa-miR-6800-5p</u> |
| PNPLA3 | 83 | <u>hsa-miR-3171</u> |
| PNPLA3 | 82 | <u>hsa-miR-8064</u> |
| PNPLA3 | 81 | <u>hsa-miR-1304-3p</u> |
| PNPLA3 | 81 | <u>hsa-miR-8075</u> |
| MBOAT7 | 96 | <u>hsa-miR-4700-5p</u> |
| MBOAT7 | 94 | <u>hsa-miR-24-3p</u> |
| MBOAT7 | 92 | <u>hsa-miR-23a-3p</u> |
| MBOAT7 | 92 | <u>hsa-miR-23b-3p</u> |
| MBOAT7 | 92 | <u>hsa-miR-23c</u> |
| MBOAT7 | 91 | <u>hsa-miR-6739-5p</u> |
| MBOAT7 | 91 | <u>hsa-miR-8089</u> |
| MBOAT7 | 91 | <u>hsa-miR-6733-5p</u> |
| MBOAT7 | 91 | <u>hsa-miR-4667-5p</u> |
| MBOAT7 | 89 | <u>hsa-miR-450a-1-3p</u> |
| MBOAT7 | 89 | <u>hsa-miR-3153</u> |
| MBOAT7 | 84 | <u>hsa-miR-4725-3p</u> |
| MBOAT7 | 82 | <u>hsa-miR-4731-5p</u> |
| MBOAT7 | 81 | <u>hsa-miR-610</u> |
| IRS1 | 100 | <u>hsa-miR-5011-5p</u> |
| IRS1 | 100 | <u>hsa-miR-3148</u> |
| IRS1 | 100 | <u>hsa-miR-8485</u> |
| IRS1 | 100 | <u>hsa-miR-190a-3p</u> |
| IRS1 | 99 | <u>hsa-miR-5696</u> |
| IRS1 | 98 | <u>hsa-miR-922</u> |
| IRS1 | 97 | <u>hsa-miR-3646</u> |
| IRS1 | 97 | <u>hsa-miR-660-5p</u> |
| IRS1 | 97 | <u>hsa-miR-6071</u> |
| IRS1 | 97 | <u>hsa-miR-651-3p</u> |
| IRS1 | 97 | <u>hsa-miR-544b</u> |
| IRS1 | 97 | <u>hsa-miR-4291</u> |
| IRS1 | 96 | <u>hsa-miR-3619-5p</u> |
| IRS1 | 96 | <u>hsa-miR-128-3p</u> |
| IRS1 | 96 | <u>hsa-miR-3919</u> |
| IRS1 | 96 | <u>hsa-miR-1284</u> |
| IRS1 | 95 | <u>hsa-miR-216a-3p</u> |
| IRS1 | 95 | <u>hsa-miR-761</u> |
| IRS1 | 95 | <u>hsa-miR-7-5p</u> |
| IRS1 | 95 | <u>hsa-miR-214-3p</u> |
| IRS1 | 94 | <u>hsa-miR-4503</u> |
| IRS1 | 94 | <u>hsa-miR-4324</u> |
| IRS1 | 94 | <u>hsa-miR-4789-3p</u> |
| IRS1 | 93 | <u>hsa-miR-570-3p</u> |
| IRS1 | 93 | <u>hsa-miR-3681-3p</u> |
| IRS1 | 92 | <u>hsa-miR-12122</u> |
| IRS1 | 92 | <u>hsa-miR-7160-5p</u> |
| IRS1 | 92 | <u>hsa-miR-5692a</u> |
| IRS1 | 91 | <u>hsa-miR-4799-5p</u> |
| IRS1 | 90 | <u>hsa-miR-6828-3p</u> |
| IRS1 | 90 | <u>hsa-miR-3941</u> |
| IRS1 | 90 | <u>hsa-miR-4670-3p</u> |
| IRS1 | 89 | <u>hsa-miR-654-5p</u> |
| IRS1 | 89 | <u>hsa-miR-3149</u> |
| IRS1 | 89 | <u>hsa-miR-541-3p</u> |
| IRS1 | 89 | <u>hsa-miR-96-5p</u> |
| IRS1 | 88 | <u>hsa-miR-6739-3p</u> |
| IRS1 | 88 | <u>hsa-miR-1250-3p</u> |
| IRS1 | 88 | <u>hsa-miR-181c-3p</u> |
| IRS1 | 88 | <u>hsa-miR-3180-5p</u> |
| IRS1 | 88 | <u>hsa-miR-5093</u> |
| IRS1 | 87 | <u>hsa-miR-30b-5p</u> |
| IRS1 | 87 | <u>hsa-miR-30e-5p</u> |
| IRS1 | 87 | <u>hsa-miR-30c-5p</u> |
| IRS1 | 87 | <u>hsa-miR-466</u> |
| IRS1 | 87 | <u>hsa-miR-30a-5p</u> |
| IRS1 | 87 | <u>hsa-miR-30d-5p</u> |
| IRS1 | 86 | <u>hsa-miR-3617-3p</u> |
| IRS1 | 86 | <u>hsa-miR-1271-5p</u> |
| IRS1 | 86 | <u>hsa-miR-4328</u> |
| IRS1 | 86 | <u>hsa-miR-12133</u> |
| IRS1 | 86 | <u>hsa-miR-4729</u> |
| IRS1 | 85 | <u>hsa-miR-1245b-3p</u> |
| IRS1 | 85 | <u>hsa-miR-6124</u> |
| IRS1 | 85 | <u>hsa-miR-12120</u> |
| IRS1 | 85 | <u>hsa-miR-3613-3p</u> |
| IRS1 | 85 | <u>hsa-miR-3928-3p</u> |
| IRS1 | 85 | <u>hsa-miR-1323</u> |
| IRS1 | 85 | <u>hsa-miR-183-5p</u> |
| IRS1 | 85 | <u>hsa-miR-1200</u> |
| IRS1 | 84 | <u>hsa-miR-4666a-3p</u> |
| IRS1 | 84 | <u>hsa-miR-7152-5p</u> |
| IRS1 | 84 | <u>hsa-miR-4680-3p</u> |
| IRS1 | 84 | <u>hsa-miR-550a-3p</u> |
| IRS1 | 82 | <u>hsa-miR-548o-3p</u> |
| IRS1 | 82 | <u>hsa-miR-7162-3p</u> |
| IRS1 | 81 | <u>hsa-miR-302b-5p</u> |
| IRS1 | 81 | <u>hsa-miR-32-3p</u> |
| IRS1 | 81 | <u>hsa-miR-6759-3p</u> |
| IRS1 | 81 | <u>hsa-miR-3133</u> |
| IRS1 | 81 | <u>hsa-miR-2115-5p</u> |
| IRS1 | 81 | <u>hsa-miR-302d-5p</u> |
| IRS1 | 80 | <u>hsa-miR-7856-5p</u> |
| IRS1 | 80 | <u>hsa-miR-559</u> |
| IRS1 | 80 | <u>hsa-miR-890</u> |
| IRS1 | 80 | <u>hsa-miR-1261</u> |
| SLC2A1 | 96 | <u>hsa-miR-5011-5p</u> |
| SLC2A1 | 93 | <u>hsa-miR-4427</u> |
| SLC2A1 | 92 | <u>hsa-miR-4758-5p</u> |
| SLC2A1 | 92 | <u>hsa-miR-1238-5p</u> |
| SLC2A1 | 91 | <u>hsa-miR-92a-1-5p</u> |
| SLC2A1 | 90 | <u>hsa-miR-6077</u> |
| SLC2A1 | 88 | <u>hsa-miR-152-3p</u> |
| SLC2A1 | 88 | <u>hsa-miR-148a-3p</u> |
| SLC2A1 | 88 | <u>hsa-miR-6081</u> |
| SLC2A1 | 88 | <u>hsa-miR-148b-3p</u> |
| SLC2A1 | 87 | <u>hsa-miR-4653-3p</u> |
| SLC2A1 | 86 | <u>hsa-miR-4742-3p</u> |
| SLC2A1 | 86 | <u>hsa-miR-8055</u> |
| SLC2A1 | 86 | <u>hsa-miR-3664-3p</u> |
| SLC2A1 | 84 | <u>hsa-miR-1263</u> |
| SLC2A1 | 84 | <u>hsa-miR-223-5p</u> |
| SLC2A1 | 81 | <u>hsa-miR-1266-5p</u> |
| SLC2A1 | 81 | <u>hsa-miR-1284</u> |
| SLC2A1 | 81 | <u>hsa-miR-6773-5p</u> |
| SLC2A1 | 81 | <u>hsa-miR-4518</u> |
| SLC2A1 | 80 | <u>hsa-miR-3163</u> |
| SLC2A1 | 80 | <u>hsa-miR-8063</u> |
| SLC2A1 | 80 | <u>hsa-miR-3140-3p</u> |
| SLC2A1 | 80 | <u>hsa-miR-548t-3p</u> |
| SLC2A1 | 80 | <u>hsa-miR-548aa</u> |
| SLC2A1 | 80 | <u>hsa-miR-548ap-3p</u> |
| SLC2A1 | 80 | <u>hsa-miR-140-5p</u> |
| SLC2A1 | 80 | <u>hsa-miR-1277-5p</u> |
| TM6SF2 | 87 | <u>hsa-miR-6778-5p</u> |
| TM6SF2 | 87 | <u>hsa-miR-1233-5p</u> |
| TM6SF2 | 85 | <u>hsa-miR-488-3p</u> |
| TM6SF2 | 85 | <u>hsa-miR-5681a</u> |
| TMC4 | 91 | <u>hsa-miR-11400</u> |

### DIANA-microT analysis

Continuing with our miRNA elucidation process, further analysis of the candidate genes was carried out with DIANA (DNA Intelligent Analysis) Lab miRNA target prediction tool DIANA-microT 5.0. ^26^ This algorithm works on parameters that are calculated individually for each miRNA and for each miRNA recognition element or MRE and takes into account binding characteristics and conservation levels. The miRNA: target interaction is primarily judged on the basis of the total score or the miRNA targeted gene (miTG) score and this predicted score is the sum of the conserved and non-conserved MREs of a particular gene. Along-with this, a signal-to-noise ratio or SNR and a precision score for each interaction is also provided for estimating the confidence of the predicted result. The DIANA microT web server allows one to search miRNA targets on a specific gene and using this feature the candidate genes were searched one by one for prospective miRNAs targeting them using a score threshold of 7.0. This web server apart from the main parameters mentioned above also gives information about the binding type, UTR position, conservation and online linkages to bibliographical and biological resources for further analysis. A list of the DIANA microT 5.0 analysis of the candidate genes with significance values (above 0.90) and their miRNAs are given in Table 3 of this study.

**Table 3:**
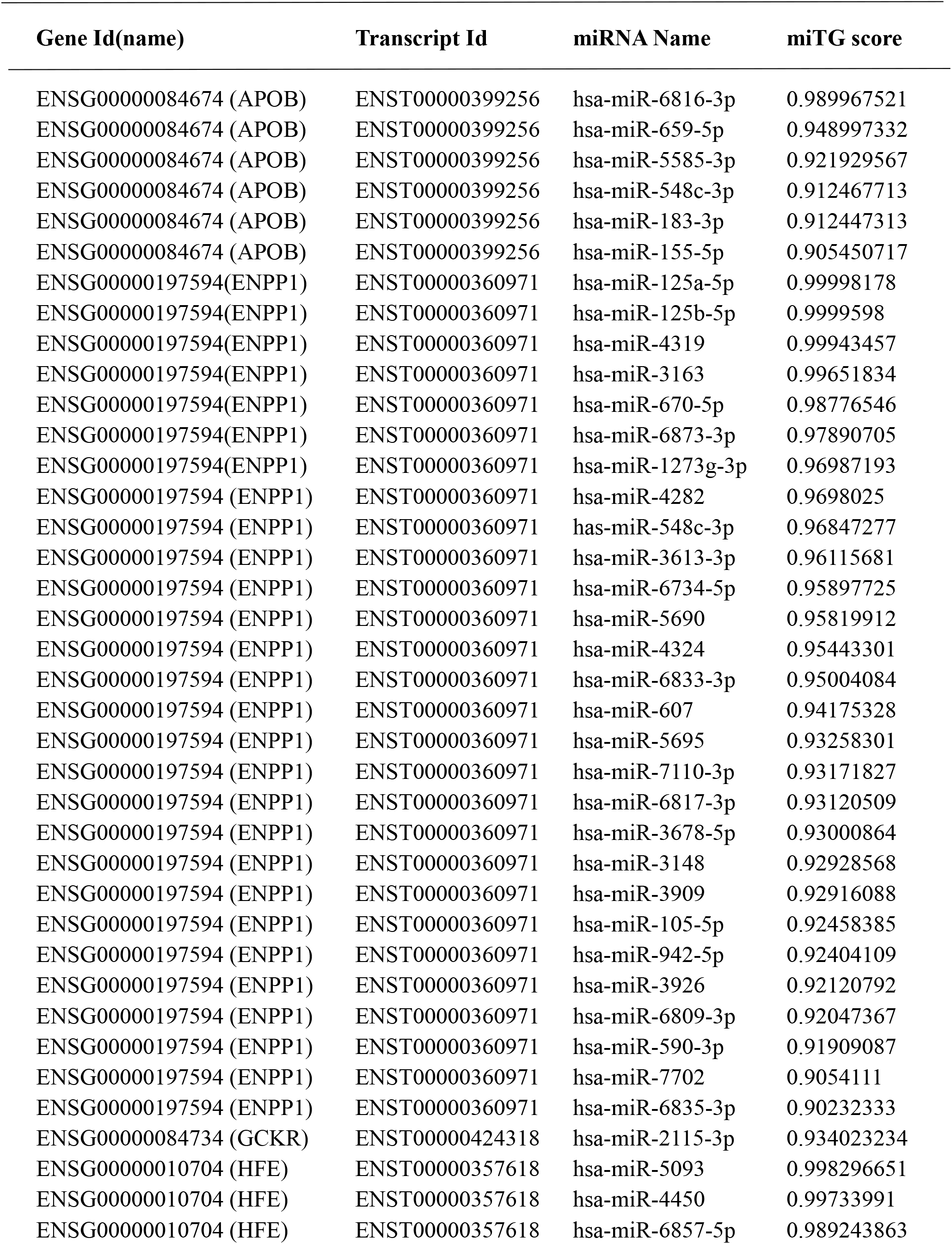
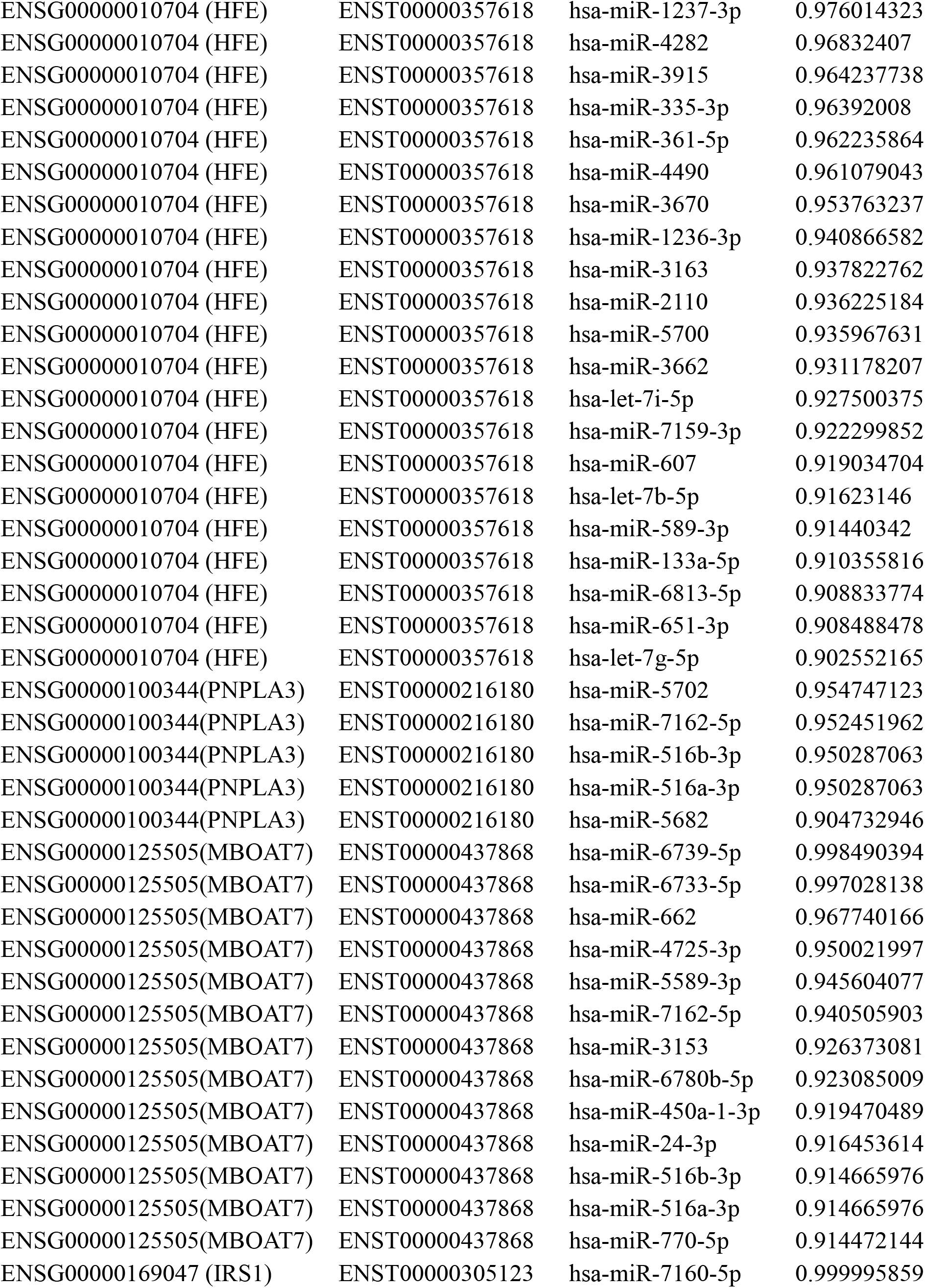
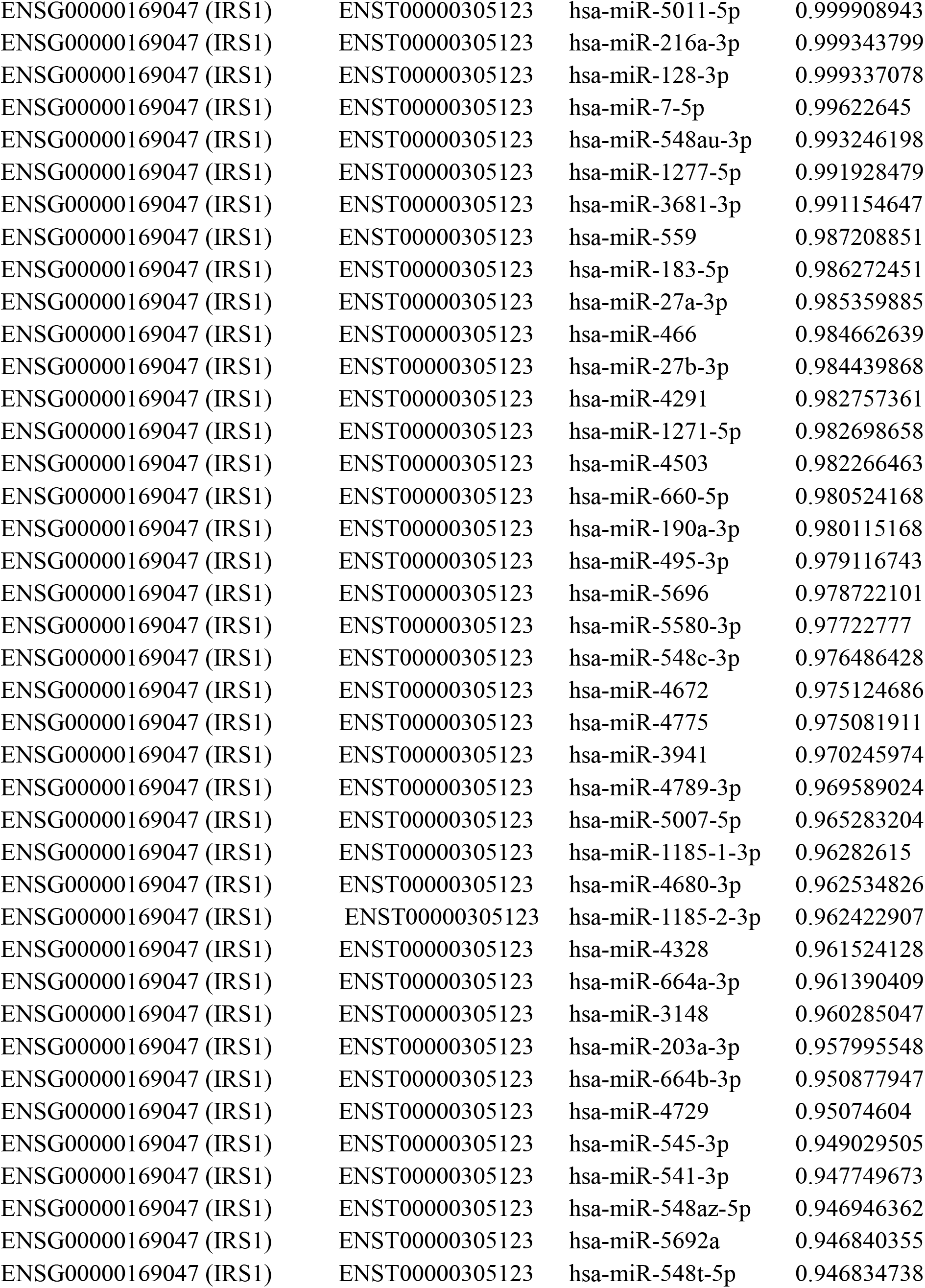
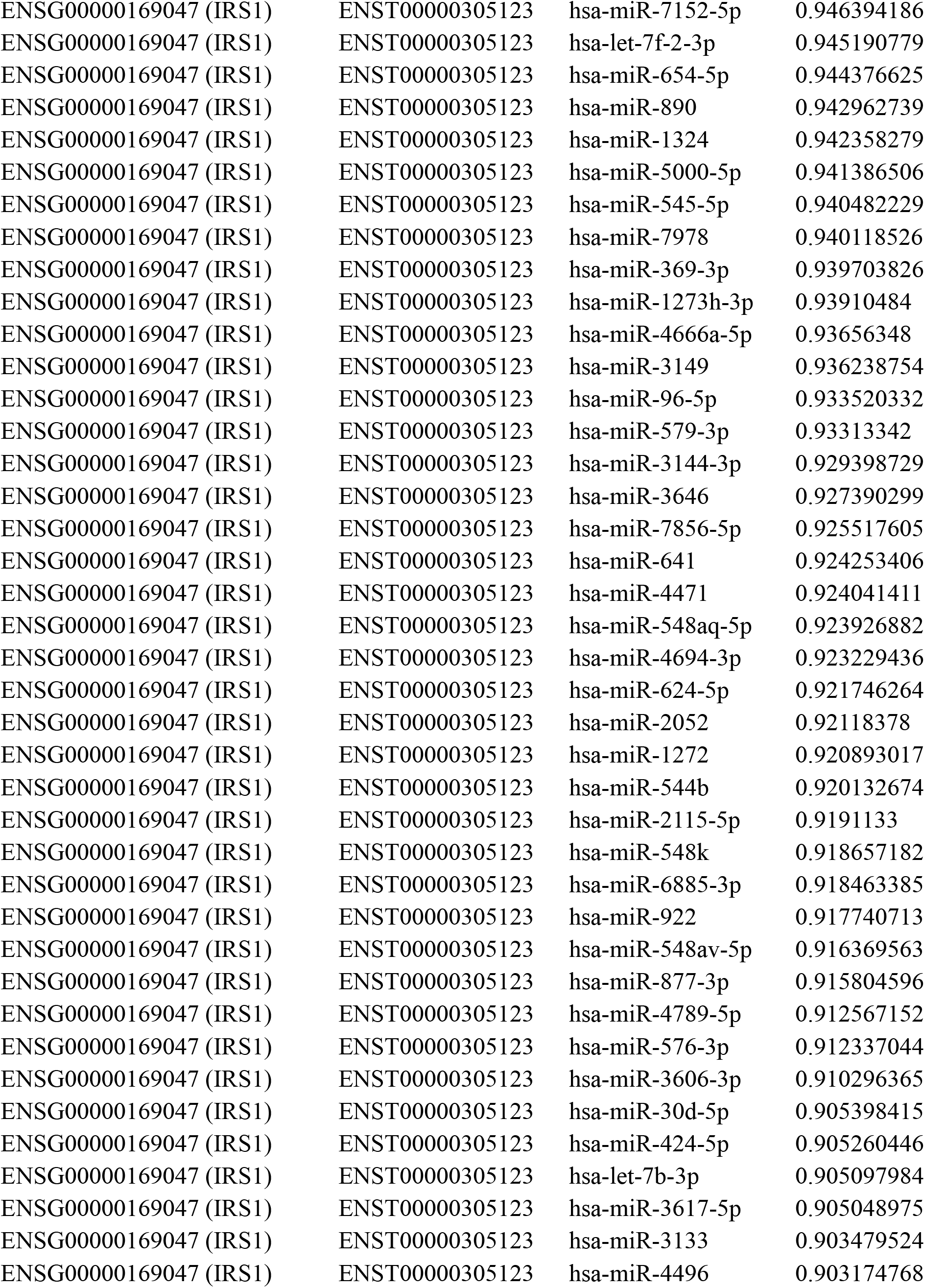
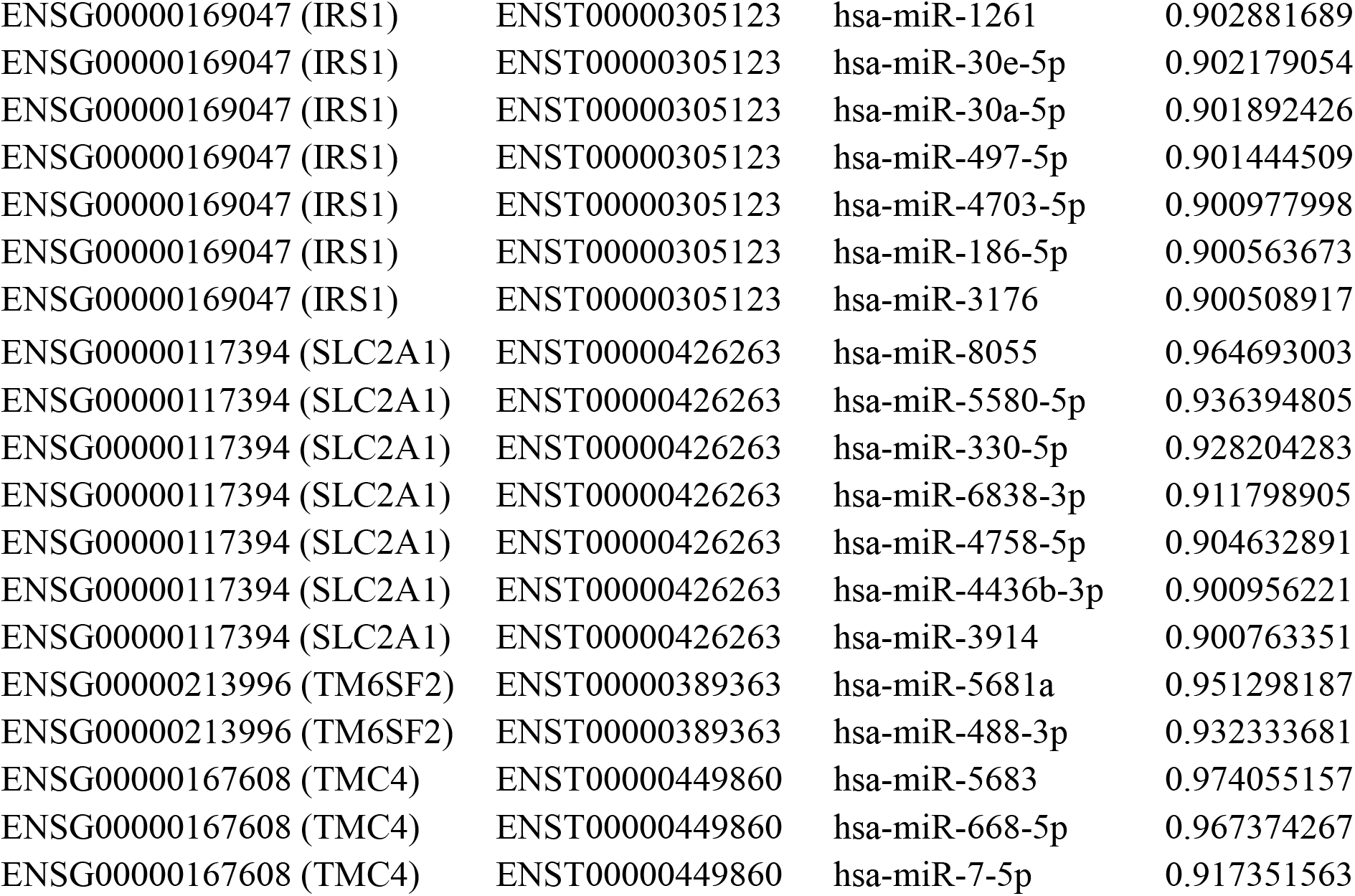
The above table shows candidate genes with miRNAs targeting them along-with significant miTG scores (≥ 0.90) from DIANA microT analysis.

### miRTarBase analysis for experimentally validated miRNAs

In order to validate our search strategy, the same gene dataset was used for identification of miRNAs using a database called miRTarBase.^27^ miTarBase is a comprehensive database containing collections of experimentally validated miRNA-Target interactions or MTIs. These MTIs are collected manually from relevant literature reporting functional miRNA studies. The experimental validation of these miRNAs is mostly done using reporter assays, western blots or microarray experiments with over-expression or knockdown of miRNAs. miRTarBase contains more than 422517 experimentally validated MTIs involving 4076 miRNAs and 23,054 target genes. Using this database, we investigated whether for any of our candidate gene(s), there is experimental evidence of miRNA-Target Interactions from literature using the validation methods. For this purpose, all the candidate genes were searched in miRTarBase by using the ‘Gene Symbol’ option and ‘Human’ in the species option. One can also browse through the gene list provided under a specific species like Human and check whether one’s gene of interest is listed in the database. When a specific hit for a target gene is found, information about both the pre- and mature miRNA including the secondary structure as well as the mature sequence is provided along-with information about the target gene, the validation methods used and the relevant literature report. Using miRTarBase, all the 10 genes were selected in the database according to the given protocol and the result of this study is given in Table 4.

**Table 4:** Candidate genes showing experimentally validated miRNA-target interaction in miRTarBase and showing validation methods with only strong evidence w.r.t experimental methodology.

|  |  |  |  |  | Validation methods |  |  |
| --- | --- | --- | --- | --- | --- | --- | --- |
|  |  |  |  |  | Strong | Evidence |  |
| ID | Species<br>(miRNA) | Species<br>(Target) | miRNA | Target | Reporter gene | Western blot | qPCR |
| MIRT731658 | Homo sapiens | Homo sapiens | hsa-miR-548p | APOB | + | + | + |
| MIRT003807 | Homo sapiens | Homo sapiens | hsa-miR-7-5p | IRS1 |  | + |  |
| MIRT000731 | Homo sapiens | Homo sapiens | hsa-miR-145-5p | IRS1 | + | + | + |
| MIRT004355 | Homo sapiens | Homo sapiens | hsa-miR-126-3p | IRS1 | + | + | + |
| MIRT006859 | Homo sapiens | Homo sapiens | hsa-miR-148a-3p | IRS1 | + | + |  |
| MIRT006860 | Homo sapiens | Homo sapiens | hsa-miR-152-3p | IRS1 | + | + |  |
| MIRT437910 | Homo sapiens | Homo sapiens | hsa-miR-105-5p | IRS1 | + | + |  |
| MIRT732116 | Homo sapiens | Homo sapiens | hsa-miR-1225-5p | IRS1 | + | + | + |
| MIRT734155 | Homo sapiens | Homo sapiens | hsa-miR-144-3p | IRS1 | + | + | + |
| MIRT735025 | Homo sapiens | Homo sapiens | hsa-miR-628-5p | IRS1 | + |  |  |
| MIRT735485 | Homo sapiens | Homo sapiens | hsa-miR-384 | IRS1 | + | + | + |
| MIRT438642 | Homo sapiens | Homo sapiens | hsa-miR-1291 | SLC2A1 | + | + | + |
| MIRT699417 | Homo sapiens | Homo sapiens | hsa-miR-22-3p | SLC2A1 | + | + | + |
| MIRT734315 | Homo sapiens | Homo sapiens | hsa-miR-148b-3p | SLC2A1 | + | + | + |
| MIRT735435 | Homo sapiens | Homo sapiens | hsa-miR-132-3p | SLC2A1 | + | + | + |
| MIRT735450 | Homo sapiens | Homo sapiens | hsa-miR-150-5p | SLC2A1 | + | + |  |
| MIRT054541 | Homo sapiens | Homo sapiens | hsa-miR-200b-3p | HFE |  | + | + |
| MIRT054542 | Homo sapiens | Homo sapiens | hsa-miR-200a-3p | HFE |  | + | + |
| MIRT054543 | Homo sapiens | Homo sapiens | hsa-miR-200c-3p | HFE |  | + | + |

Based on this *in silico* elucidation analysis, a meta-list of candidate genes and their elucidated cognate miRNAs common to both miRDB and DIANA were created and listed in Table 5. The entries here were searched to identify putative hits having experimental validation in miRTarBase database also and only those hits with strong experimental evidence selected for further discussion.

**Table 5:** Comparative data of candidate genes along-with their cognate miRNAs from all three databases used in the miRNA elucidation pipeline. miRNAs common in both miRDB and DIANA as well as having strong experimental evidence as per miRTarBase are identified in bold.

| Gene Name | mi-RDB | T-CDS | miRTarBase<br>Strong<br>evidence |
| --- | --- | --- | --- |
| APOB | **<u>hsa-miR-548p</u>** | **<u>hsa-miR-548-3p</u>** | + |
| IRS1 | **<u>hsa-miR-7-5p</u>** | **<u>hsa-miR-7-5p</u>** | + |

## Results and Discussion

The gene list (Table 1) was obtained from the study of Alessandra *et. al.,* and the genes were put through the *in silico* miRNA elucidation pipeline. In this study, miRDB and T-CDS were used to identify putative miRNA binding sites. This was followed by analysis using miRTarBase which is an experimentally validated database of miRNA-target interactions. In miRDB study, a target score of greater than 80 is selected as a statistically significant value according to the miRDB program itself. A list of these candidate genes with various miRNA’s targeting them is given in Table 2. This list contains all the genes with miRNAs having a minimum target score of 80 and above only. In T-CDS study, a target score of greater than 0.90 is selected as a statistically significant value which is according to the T-CDS program. A list of these candidate genes with various miRNA’s targeting them is given in Table 3 and this list contains all the genes having a minimum target score of 0.90 and above. miRTarBase is a database containing a collection of experimentally validated miRNA-Target interactions. This database has been used to check miRNA-Target interactions and Table 4 contains the candidate miRNAs which are validated according to miRTarBase criteria. Table 5 of this study is comparative data of candidate genes having significant miRNA hits which have been found to be common in both miRDB and T-CDS programs. The common miRNA hits of miRDB and DIANA T-CDS were also searched in miRTarBase database.

NAFLD with its abnormal fat accumulation is the most common cause of chronic liver disease especially in the western world.^28^ This disorder can manifest in multiple ways including steatohepatitis, fibrosis and cirrhosis.^29^ Additionally, NAFLD is also strongly associated with metabolic syndrome and cardiovascular disease and a subset of such patients may end up with hepatocellular carcinoma (HCC). The hallmark of NAFLD is the accumulation of triglycerides within hepatocytes with increased *de novo* lipogenesis, high adipose tissue lipolysis, obesity and diabetes.^29^ The progression of NAFLD to fibrosis is mediated by a complex web of factors including inflammatory mediators, lipotoxicity, fatty acids, dietary factors, genetic factors and epigenetics.^30^ However, despite understanding the factors responsible for NAFLD and its progression, little is known about the regulation of these factors at both transcriptional or post-transcriptional levels as well as the identity of novel biomarkers that can help in predicting clinical outcome. Therefore, in this study, we have attempted to elucidate the novel regulatory miRNA candidates involved in post-transcriptional regulation of cognate genes implicated in the pathophysiology of NAFLD and its progression using an *in silico* approach.

miRNAs as endogenous single-stranded RNAs have emerged as very important component of gene expression regulation through post-transcriptional mechanisms in wide variety of pathological conditions.^31^ The stability of miRNA in circulatory system makes them an attractive candidate for biomarker discovery.^32^ The input genes in our study were taken from the review of Alessandra *et. al.*, and in that they were classified under genetic factors category involved in NAFLD/NASH pathophysiology. The various genes belong to diverse physiological domains and all have been found to be associated significantly in genetic studies in NAFLD cases in multiple population types. The functional categories in this subset of genes include those involved in retinol metabolism, synthesis of fatty acids, lipid transport, oxidative stress, glucose metabolism and fibrosis.

miRNA have a very important role in metabolic homeostasis in an individual. In the liver, miR-112 influences the genes responsible for the metabolism of hepatic cholesterol and lipids. Using antisense methods, obstruction of miR-112 resulted in the reduction of plasma cholesterol levels in mice and chimpanzees.^33,34^ A separate study showed that in a mice model where miR-122 encoding gene was deleted, development of steatohepatitis, HCC and fibrosis was observed.^35^ For normal liver homeostasis, miR-122 is essential and the decreased levels of miR-122 have damaging effects on the liver. ^35^ Lipid and cholesterol regulatory genes are regulated by miRNA such as miR-33, miR-34, miR-103, miR-104 and miR-370.^36^

The mitochondrial enzyme, carnitine palmitoyl transferase that is involved in the movement of long-chain fatty acids across the membrane is affected by miR-370.^37^ At many levels miRNAs are responsible for the regulation of liver fibrosis. Upon fibrogenic injury of the liver, the Hepatic Stellate Cells (HSCs), a primary type of cells responsible for liver fibrosis, experience growth and differentiation into myofibroblast-like cells. Many varieties of miRNA are identified in regulating the HSC stimulation.^14^ Increased serum levels of miR-571 are considered as a potential biomarker for liver fibrosis.^38^ In bile-duct-ligated rats, the decrease in miRNA-150 and miRNA-194 has been identified. This miRNA-150 and miRNA-194 targeted c-Myc and Rac-1 respectively, thus restraining the HSC activation.^39^

Apart from the above abnormalities, increased levels of miR-705, miR-1224, miR-486, miR-320 and decreased levels of some other miRNAs like miR-192, miR-183, miR-199a, miR27b, miR-214 was observed in the livers of the mice subjected to chronic feeding of alcohol.^40^ In alcoholic steatohepatitis and NASH, the Kuffer cell activation is common. Up-regulation of miRNA-155 occurs in Kuffer cell in ALD, so we can theorize that it is the same for NASH as well. In patients with NASH, altered hepatic expression has been found.^41^

In Alcoholic Steatohepatitis and NASH, the Kupffer cell activation is common. Up-regulation of miRNA-155 occurs in Kupffer cells in ALD, so we can theorize that it is the same for NASH as well. In patients with NASH, altered hepatic expression has been found.^41^ It has been seen that, in patients with NASH miRNA-122 decreases as their disease progresses. These decreased levels of miRNA-122 contribute to the metabolism of altered hepatic lipids.^36^

In our *in silico* study, Table 5 shows the comparison of DIANA-microT, miRDB and miRTarBase analysis and in this table miR-7-5p (miR-7 family) is present and this have been found in literature to be an anti-oncogenic miRNA and its repression by a complex mechanism causes manifestation of HCC which can be observed is almost 22% of NAFLD cases in western countries.^7,42^ In the same table, miR-548p and 548-3p (miR-548 family) are also present in all three above mentioned tools. miR-548 is a large, poorly conserved primate-specific miRNA family and it has been implicated in multiple pathological processes like cancer and signalling pathways.^43^ miR-548a-5p been reported to negatively regulate the tumour inhibitor gene Tg737 and promote tumorigenesis *in vitro* and *in vivo* especially with respect to HCC cell proliferation and apoptosis. ^44^

From the above discussion, it is evident that certain miRNA’s like miR-7-5p and miR548p/548-3p elucidated in our study are involved in pathophysiology of NAFLD/NASH. Such miRNA candidates can also serve as prognostic markers for accurate assessment of NAFLD/NASH early on. Our analysis however has its own limitation as far as the providing functional significance and information is concerned. This analysis gives an idea of specific miRNA targeting gene implicated in liver disease. Our study also does not exclude the involvement of other miRNA(s) targeting such pathways. This analysis is putative and functional aspects of this post-transcriptional regulation needs to be validated experimentally. Some techniques by which such validation in clinical sample can be achieved includes Reporter Assay, qPCR, and pSILAC etc.

This study represents an advance in biomedical science because it provides a novel and unique approach to understand the complex regulation of genes involved in NAFLD/NASH as well as provide prognostic /diagnostic markers for this disease.

## Acknowledgements

We would like to acknowledge the support received from Dr. Kiran Mangaonkar, Principal, G.N. Khalsa College of Arts, Science & Commerce (Autonomous), Mumbai. We would also like to thank Dr. Jaimini Sarkar for her help and support in editing this paper.

## Conflict of Interest

The authors report No Conflict of Interest.

## Funding details

Not Applicable

## References

1. Gregory A. Michelotti, Mariana V. Machado and Anna Mae Diehl (2013) ‘NAFLD, NASH and Liver Cancer’, Nature Reviews Gastroenterology & Hepatology, vol 10, pp.656–665.

2. Mariana Lazo, Rubenhernaez, Mark S. Eberhardt, Susanne Bonekamp, Ihab Kamel, Eliseo Guallar, Ayman Koteish, Frederick L. Brancati, And Jeanne M. Clark (2012) ‘Prevalence of Nonalcoholic Fatty Liver Disease In The United States: The Third National Health And Nutrition Examination Survey, 1988–1994’, American Journal Of Epidemiology, Vol. 178, No.1, pp. 38–45.

3. Ankita Chatterjee, Analabha Basu, Abhijit Chowdhury, Kausik Das, Neeta Sarkar-Roy, Partha P. Majumder, Priyadarshi Basu (2015) ‘Comparative analyses of genetic risk prediction methods reveal extreme diversity of genetic predisposition to non-alcoholic fatty liver disease (NAFLD) among ethnic populations of India’, Journal of Genetics, Vol. 94 No. 1, pp. 105–112.

4. Jeanne M. Clark, M.D., M.P.H., Frederick L. Brancati, M.D., M.H.S., And Anna Mae Diehl, M.D (2003) ‘The Prevalence and Etiology Of Elevated Aminotransferase Levels in The United States’, The American Journal of Gastroenterology, Vol. 98, No. 5, pp. 960–967.

5. Arthur J. Mccullough, (2013) ‘The Clinical Features, Diagnosis and Natural History of Non-alcoholic Fatty Liver Disease’, Clinics in Liver Disease, Vol. 8, Pp 521–533.

6. Kohichiroh Yasui, Etsuko Hashimoto, Katsutoshi Tokushige, Kazuhiko Koike, Toshihide Shima, Yoshihiro Kanbara, Toshiji Saibara, Hirofumi Uto, Shiro Takami, Miwa Kawanaka, Yasuji Komo (2012) ‘Clinical And Pathological Progression Of Non-Alcoholic Steatohepatitis To Hepatocellular Carcinoma’, The Japan Society Of Hepatology, Vol. 368 No.20, pp.1–7.

7. Ju Dong Yang, Bohyun Kim, Schuyler O. Sanderson, Jennifer L. St. Sauver, Barbara P. Yawn, Rachel A. Pedersen, BA; Joseph J. Larson, BA; Terry M. Therneau, Lewis R. Roberts, W. Ray Kim (2012) ‘Hepatocellular Carcinoma in Olmsted County, Minnesota, 1976-2008’, Mayo Clin Proc, Vol. 87 No.1, pp.9–16.

8. Anna Ludovica Fracanzani, Luca Valenti, Elisabetta Bugianesi, Marco Andreoletti, Agostino Colli, Ester Vanni, Cristina Bertelli, Erika Fatta, Daniela Bignamini, Giulio Marchesini, Silvia Fargion (2008) ‘Risk of Severe Liver Disease in Non-alcoholic Fatty Liver Disease with Normal Aminotransferase Levels: A Role for Insulin Resistance and Diabetes’, Hepatology, Vol. 48 No. 3, pp.792–798.

9. V.W.S. Wong, G.L.H. Wong, S.W.C. Tsang, A.Y. Hui, A.W.H. Chan, P.C.L. Choi, A.M.L. Chim, S. Chu, F.K.L. Chan, J. J.Y. Sung, H. L. Y. Chan (2008) ‘Metabolic And Histological Features Of Non-alcoholic Fatty Liver Disease Patients With Different Serum Alanine Aminotransferase Levels’, Aliment Pharmacol and Therapeutics, 2008, Vol. 29, pp. 387–396.

10. Onpan Cheung and Arun J. Sanyal (2009), ‘Recent Advances in Non-alcoholic Fatty Liver Disease’, Current Opinion in Gastroenterology, Vol. 25, pp 230–237.

11. Natsuko Kawada, Kazuho Imanaka, Tsukasa Kawaguchi, Chie Tamai, Ryu Ishihara, Takashi Matsunaga, Kunihito Gotoh, Terumasa Yamada, Yasuhiko Tomita (2009) ‘Hepatocellular Carcinoma Arising from Non-Cirrhotic Non-alcoholic Steatohepatitis’, J Gastroenterol, Vol. 44, pp 1190–1194.

12. Katsutoshi Tokushige, Etsuko Hashimoto, Yoshinori Horie, Makiko Taniai, Susumu Higuch (2011) ‘Hepatocellular Carcinoma In Japanese Patients With Non-alcoholic Fatty Liver Disease, Alcoholic Liver Disease, And Chronic Liver Disease Of Unknown Etiology: Report Of The Nationwide Survey’, J Gastroenterol, Vol. 46, pp 1230–1237.

13. Lee RC, Feinbaum RL, Ambros V. (1993) ‘The C. elegans heterochronic gene lin-4 encodes small RNAs with antisense complementarity to lin-14’, Cell, Vol.75 No. 5, 843–854.

14. Carlos J Pirola, Tomas Fernández Gianotti, Gustavo O Castano, Pablo Mallardi, Julio San Martino, María Mora Gonzalez Lopez Ledesma, Diego Flichman, Faridodin Mirshahi, Arun J Sanyal, and Silvia Sookoian (2015) ‘Circulating Microrna Signature in Non-Alcoholic Fatty Liver Disease: From Serum Non-Coding Rnas To Liver Histology and Disease Pathogenesis’, Gut, Vol. 65 No. 5, pp. 800–812.

15. Gyongyi Szabo and Shashi Bala (2013) ‘MicroRNAs In Liver Disease’, Gastroenterol Hepatol, Vol 10 No. 9, pp 542–552.

16. Alessandra Caligiuri, Alessandra Gentilini and Fabio Marra (2016) ‘Molecular Pathogenesis of NASH’, International Journal of Molecular Science, Vol. 17, Pp 1–34.

17. Leigh-Ann Macfarlane and Paul R. Murphy (2010) ‘MicroRNA: Biogenesis, Function and Role in Cancer’, Current Genomics, Vol 11 No. 7, pp 537–561.

18. S.H. Hsu, B. Wang, J. Kota, J. Yu, S. Costinean, H. Kutay, L. Yu, S. Bai, K. La Perle, R.R. Chivukula, H. Mao, M. Wei, K.R. Clark, J.R. Mendell, M.A. Caligiuri, S.T. Jacob, J.T. Mendell, K. Ghoshal (2012) ‘Essential metabolic, anti-inflammatory, and anti-tumorigenic functions of miR-122 in liver’, J. Clin. Invest. Vol. 122, pp. 2871–2883.

19. Y. Xu, M. Zalzala, J. Xu, Y. Li, L. Yin, Y. Zhang (2015) ‘A metabolic stress-inducible miR-34a-HNF4alpha pathway regulates lipid and lipoprotein metabolism’, Nat. Commun. Vol. 6, pp. 7466.

20. Q. Xu, Y. Li, Y.F. Shang, H.L. Wang, M.X. Yao (2015), ‘miRNA-103: molecular link between insulin resistance and non-alcoholic fatty liver disease’, World J. Gastroenterol Vol. 21, pp. 511–516.

21. X. Pan, P. Wang, J. Luo, Z. Wang, Y. Song, J. Ye, X. Hou (2015) ‘Adipogenic changes of hepatocytes in a high-fat diet-induced fatty liver mice model and non-alcoholic fatty liver disease patients’, Endocrine, Vol. 48, pp 834–847.

22. M. Trajkovski, J. Hausser, J. Soutschek, B. Bhat, A. Akin, M. Zavolan, M.H. Heim, M. Stoffel (2011), ‘MicroRNAs 103 and 107 regulate insulin sensitivity’, Nature Vol. 474, pp. 649–653.

23. A.M. Miller, D.S. Gilchrist, J. Nijjar, E. Araldi, C.M. Ramirez, C.A. Lavery, C. Fernandez-Hernando, I.B. McInnes, M. Kurowska-Stolarska (2013) ‘MiR-155 has a protective role in the development of non-alcoholic hepatosteatosis in mice’, PLoS One 8 e72324.

24. A.N. Mattis, G. Song, K. Hitchner, R.Y. Kim, A.Y. Lee, A.D. Sharma, Y. Malato, M.T. McManus, C.C. Esau, E. Koller, S. Koliwad, L.P. Lim, J.J. Maher, R.L. Raffai, H. Willenbring (2015), A screen in mice uncovers repression of lipoprotein lipase by microRNA-29a as a mechanism for lipid distribution away from the liver’, Hepatology Vol. 6, pp. 141–152.

25. Nathan Wong and Xiaowei Wang (2015) ‘miRDB: an online resource for microRNA target prediction and functional annotations’, Nucleic Acids Research, Vol. 43, pp 146–152.

26. M. Maragkakis, M. Reczko, V. A. Simossis, P. Alexiou, G. L. Papadopoulos, T. Dalamagas, G. Giannopoulos, G. Goumas, E. Koukis, K. Kourtis, T. Vergoulis, N. Koziris, T. Sellis, P. Tsanakas, A. G. Hatzigeorgiou (2009) ‘DIANA-microT web server: elucidating microRNA functions through target prediction’, Nucleic Acids Research, Vol. 37, pp 273–276.

27. Sheng-Da Hsu, Feng-Mao Lin, Wei-Yun Wu, Chao Liang, Wei-Chih Huang, Wen-Ling Chan, Wen-Ting Tsai, Goun-Zhou Chen, Chia-Jung Lee, Chih-Min Chiu, Chia-Hung Chien, Ming-Chia Wu, Chi-Ying Huang, Ann-Ping Tsou and Hsien-Da Huang (2011) ‘miRTarBase: a database curates experimentally validated microRNA–target interactions’, Nucleic Acids Research, Vol. 39, pp 163–169.

28. Ogawa Y, Imajo K, Yoneda M, Nakajima A (2017), ‘Update: epidemiology and pathophysiology of nonalcoholic fatty liver disease’, Nihon Shokakibyo Gakkai Zasshi Vol. 111, pp 14–24.

29. Wonhee Hur, Joon Ho Lee, Sung Woo Kim, Jung-Hee Kim, Si Hyun Bae, Minhyung Kim, Daehee Hwang, Young Seok Kim, Taesun Park, Soo-Jong Um, Byoung-Joon Song, Seung Kew Yoon (2015) ‘Downregulation of microRNA-451 in non-alcoholic steatohepatitis inhibits fatty acid-induced proinflammatory cytokine production through the AMPK/AKT pathway’, The International Journal of Biochemistry & Cell Biology, Vol. 64, pp 265–276.

30. Qiaozhu Su, Virender Kumar, Neetu Sud, Ram I. Mahato (2018) ‘Role of MicroRNAs in the pathogenesis and treatment of progressive liver injury in NAFLD and liver fibrosis’, Advanced Drug Delivery Reviews, vol 129, pp 1–26.

31. Filipowicz W, Bhattacharyya SN & Sonenberg N (2008). ‘Mechanisms of post-transcriptional regulation by microRNAs: are the answers in sight?’, Nat. Rev. Genet. Vol. 9, pp. 102–114.

32. Gyongyi Szabo, Timea Csak (2016), ‘Role of MicroRNAs in NAFLD/ NASH’, Dig Dis Sci, Vol. 61, pp. 1314–1324.

33. Jan Kru Tzfeldt, Nikolaus Rajewsky, Ravi Braich, Kallanthottathil G. Rajeev, Thomas Tuschl, Muthiah Manoharan4 & Markus Stoffel (2005) ‘Silencing of MicroRNAs In Vivo With ‘Antagomirs’, Nature, Vol 438 No. 1, pp 685–689.

34. Christine Esau, Scott Davis, Susan F. Murray, Xing Xian Yu, Sanjay K. Pandey, Michael Pear, Lynnetta Watts, Sheri L. Booten, Mark Graham, Robert McKay, Amuthakannan Subramaniam, Stephanie Propp, Bridget A. Lollo, Susan Freier, C. Frank Bennett, Sanjay Bhanot, And Brett P. Monia1 (2006), ‘Mir-122 Regulation of Lipid Metabolism Revealed By In Vivo Antisense Targeting’, Cell Metabolism, Vol. 3, pp 87–98.

35. Wei-Chih Tsai, Michael Hsiao, Ann-Ping Tsou (2012) ‘MicroRNA-122 Plays a Critical Role in Liver Homeostasis and Hepatocarcinogenesis’, The Journal of Clinical Investigation, Vol. 122 No. 8, pp 2884–2897.

36. Veerle Rottiers and Anders M. Naar (2012) ‘MicroRNAs In Metabolism and Metabolic Disorders’, Molecular Cell Biology, Vol. 13, pp 239–250.

37. Dimitrios Iliopoulos, Konstantinos Drosatos, Yaeko Hiyama, Ira J. Goldberg and Vassilis I. Zannis (2010) ‘Micro RNA-370 Control Expressions of Micro RNA-122 and Cpt1 and Affects Lipid Metabolism’, Journal of Lipid Research, Vol. 51, Pp 1513–1523.

38. Christoph Roderburg, Gerd Willem Urban, Kira Bettermann, Mihael Vucur, Henning Zimmermann, Sabine Schmidt, Jörn Janssen, Christiane Koppe, Percy Knolle, Mirco Castoldi, Frank Tacke, Christian Trautwein, (2010) ‘MicroRNA Profiling Reveals A Role Of Micro RNA In Human And Murine Liver Fibrosis’, Hepatology, Vol. 53 No. 10, pp. 209–218.

39. Senthil K. Venugopal, Joy Jiang, Tae-Hun Kim, Yong Li, Si-Si Wang, Natalie J. Torok, Jian Wu and Mark A. Zern (2009) ‘Liver Fibrosis Causes Downregulation of Mirna-150 And Mirna-194 In Hepatic Stellate Cells, and Their Overexpression Causes Decreased Stellate Cell Activation’, AJP Gastrointest Liver Physiol, Vol. 208, pp 101–106.

40. Angela Dolganiuc, Jan Petrasek, Karen Kodys, Donna Catalano, Pranoti Mandrekar, Arumugam Velayudham, And Gyongyi Szabo (2009) ‘Microrna Expression Profile in Lieber-Decarli Diet-Induced Alcoholic and Methionine Choline Deficient Diet-Induced Non-alcoholic Steatohepatitis Models in Mice’, Alcoholism: Clinical and Experimental Research, 2009, Vol. 33 No. 10, pp 1704–1710.

41. Bala S, Szabo G. (2011) ‘MicroRNA Signature in Alcoholic Liver Disease’, International Journal of Hepatology, Vol. 2012, Pp 1–6.

42. Takuma Higuchi, Hiroshi Todaka, Yasunori Sugiyama, Masafumi Ono, Nobuyuki Tamaki, Etsuro Hatano, Yuka Takezaki, Kazuhiro Hanazaki, Takeshi Miwa, Sylvia Lai, Keiko Morisawa, Masayuki Tsuda, Taketoshi Taniguchi, Shuji Sakamoto (2016) ‘Suppression of MicroRNA-7 (miR-7) Biogenesis by Nuclear Factor 90-Nuclear Factor 45 Complex (NF90-NF45) Controls Cell Proliferation in Hepatocellular Carcinoma’, Journal of Biological Chemistry, Vol. 291 No. 40, pp. 21074–21084.

43. Tingming Liang, Li Guo, Chang Liu (2012), ‘Genome-Wide Analysis of mir-548 gene Family Reveals Evolutionary and Functional Implications’, Journal of Biomedicine and Biotechnology, pp 1–8.

44. Ge Zhao, Ting Wang, Qi-Ke Huang, Meng Pu, Wei Sun, Zhuo-Chao Zhang, Rui Ling, Kai-Shan Tao (2016), ‘MicroRNA-548a-5p promotes proliferation and inhibits apoptosis in hepatocellular carcinoma cells by targeting Tg 737’, World Journal of Gasteroenterology, Vol 22, No. 23, pp 5364–5373.

